# Physical restraint induces conditioned place aversion and region-specific c-Fos activation in mice

**DOI:** 10.64898/2026.05.07.723616

**Authors:** Ella Reinders, Maya Tondravi, Se Rin Lee, Eden Beyene, Tyler Nguyen, Tara A. LeGates

**Affiliations:** Department of Biological Sciences, University of Maryland, Baltimore County (UMBC); Department of Pharmacology and Physiology, University of Maryland, School of Medicine

**Author notes:** Corresponding author: Tara LeGates, Ph.D., 1000 Hilltop Circle, Interdisciplinary Life Sciences Building 315, Baltimore, MD 21250.

**Keywords:** c-Fos, aversion, nucleus accumbens, hippocampus, conditioned place aversion

## Abstract

Linking environmental contexts with stressful experiences is critical for engaging adaptive responses necessary to avoid future threats. Yet, active context-dependent avoidance remains poorly understood. Here, we establish a restraint-induced conditioned place aversion (CPA) paradigm to examine how an acute physiological stressor acquires negative motivational value through contextual association. We found that mice repeatedly exposed to physical restraint in a contextually distinguishable chamber later avoid that location, demonstrating that restraint stress can drive learned aversion in the absence of continued exposure. To identify potential neuronal correlates underlying this learned association, we quantified c-Fos expression in several areas implicated in aversive motivation, emotional salience, and contextual encoding. We found that restraint within the context of the CPA paradigm was associated with increased c-Fos in the nucleus accumbens (NAc) and basolateral amygdala (BLA) while c-Fos expression increased in the ventral hippocampus in response to exposure to the contextual cues alone. These findings reveal region-specific activation in response to restraint stress and associated contextual cues. By connecting classical stress models and associative learning, this work provides a potential platform for further investigation of the neural mechanisms underlying stress-related negative affect and avoidance behaviors.

## 1. Introduction

Acute stress triggers adaptive responses to sharpen vigilance for threats, and associative learning links stress-inducing stimuli with the contexts in which they occur to guide future behavior [1]. These learned associations become maladaptive when they are not later extinguished or become too generalized. Such overgeneralization can trigger defensive responses to loosely related, or even unrelated stimuli, as observed in post-traumatic stress disorder (PTSD) and anxiety disorders [2].

Investigations into the neuronal correlates of this form of learning have largely focused on fear conditioning where passive defensive responses, like freezing, measure learned associations between an aversive stimulus, like footshock, and a cue or environmental context [3-6]. Active defensive responses, like avoidance, are less well studied and often involve measuring an animal’s ability to respond to a cue by moving to another part of an arena to avoid a stimulus such as footshock [7-8]. Importantly, shock delivery can occur in multiple trials if animals freeze or fail to respond in time, reinforcing the cue-stimulus association and complicating isolation of avoidance elicited by previous negatively-conditioned contextual cues alone. As avoidance often persists long after the cessation of threat, this limits our ability to relate stress-induced physiological changes to the formation of persistent stress-related memories.

Here, we examined conditioned place aversion (CPA) in response to physical restraint, a well-characterized stressor that reliably activates stress circuits and produces behavioral changes relevant to anxiety and depression [9-12]. Despite its extensive use in acute and chronic stress models, the aversive motivational properties of restraint stress have not been assessed using a contextually-based learning paradigm. Here, we show that restraint induces CPA indicating that mice can establish associations between exposure to this stressor and contextual features in the environment. We then used c-Fos expression to determine whether brain regions commonly associated with stress-related disorders might be preferentially activated during conditioning. We focused on the hippocampus (Hipp), nucleus accumbens (NAc), and basolateral amygdala (BLA), areas extensively implicated in contextual learning, motivational salience, and fear responses, respectively [13-15], and identified region-specific activation patterns.

## 2. Materials and Methods

### 2.1 Animals

Adult female and male C57BL/6J mice (17-20 weeks old) were bred in-house. Mice were group housed in a temperature- and humidity-controlled room under a 12:12-hour light-dark cycle with food and water *ad libitum*. All experiments were conducted following the regulations established by the Institutional Animal Care and Use Committee at the University of Maryland, Baltimore County.

### 2.2 Conditioned Place Aversion (CPA)

CPA took place over 5 days using an arena comprised of two chambers, distinguishable by visual cues (stripes/dots on the chamber wall), connected by a central corridor (MazeEngineers) (Figure 1a). Mice (n=12 male, 10 female) were acclimated to the testing room for 1 hour prior to the start of the experiment. *Habituation (Day 1):* Mice were allowed to freely explore the entire arena for 30 minutes. *Conditioning (Days 2-4)*: Doors were inserted to prevent mice from moving between chambers. Session 1: Mice were confined to one of the two chambers and allowed to explore the chamber for 30 minutes. Session 2: Mice were confined to the other chamber and allowed to explore for 5 minutes. They were then physically restrained in a restraint tube (internal diameter: 1”, length: 4”, IBI Scientific, Peosta, IA) for the remaining 25 minutes. Sessions 1 and 2 were conducted at least 4 hours apart. These conditioning sessions were repeated on Days 3 and 4. Session order was counterbalanced across days and animals. Chamber assignments (stripes/dots) were counterbalanced across subjects. *Testing (Day 5):* Mice were allowed to freely explore the entire arena for 20 minutes. Behavior was monitored by a camera mounted above the arena connected to a computer equipped with behavioral tracking software (ANY-maze Version 7.4). We calculated the time spent in the restraint-paired chamber as a percentage of total time spent in both chambers during habituation and testing.

**Figure 1.**
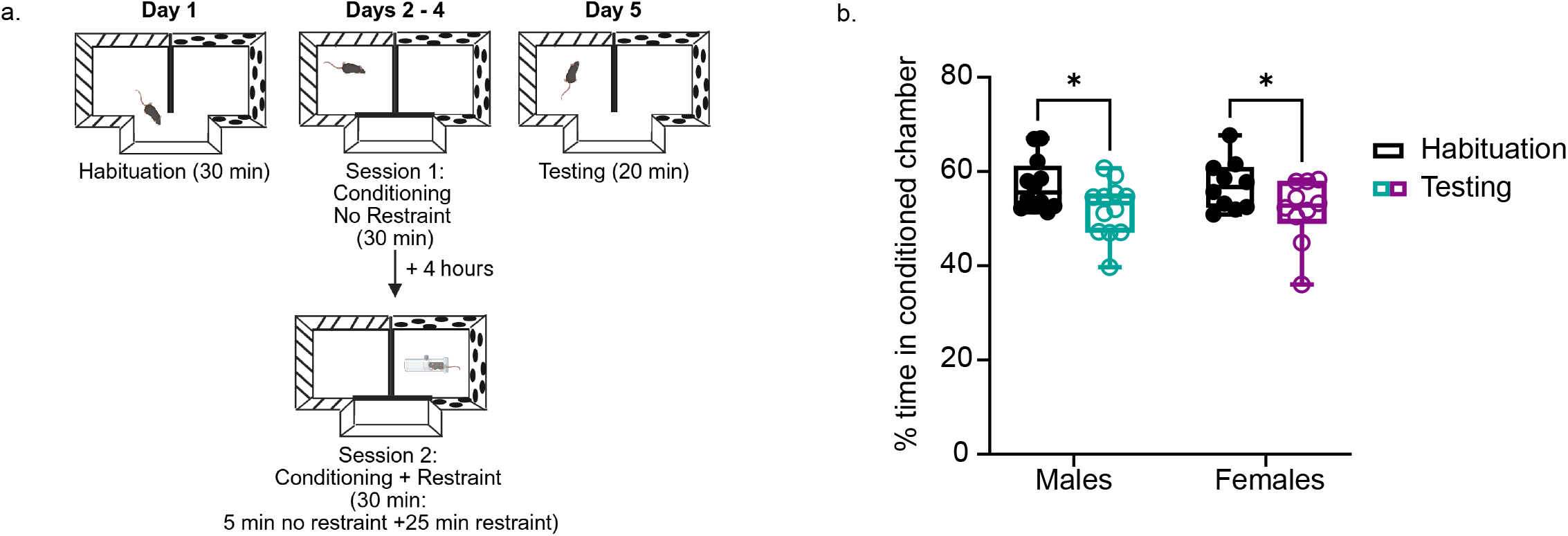
Restraint induces conditioned place aversion. a) Schematic of the CPA paradigm. b) Percent time spent in the restraint-paired chamber during habituation and testing ( n=12M, 10F; Interaction: F_(1,20)_= 0.001093, n^2^_p_ =5.463x10^-5^; Main effect of sex: F_(1,20)_= 0.007055, n^2^_p_ =0.0008; Main effect of testing stage: F_(1,20)_=14.47, n^2^_p_=0.419; *p_(male:habituation vs. testing)_=0.0222, *p_(female:habituation vs testing)_=0.0341).

### 2.3 Behavior experiment for c-Fos expression

Mice were randomly assigned to one of four groups (n=6 per group: 3 females and 3 males): (1) Home Cage No Restraint: mice remained undisturbed in their home cage, (2) Home Cage + Restraint: mice were placed in a restraint tube in their home cage for 30 minutes (3) Arena No Restraint: mice were placed in one of the CPA arena chambers and allowed to freely explore for 30 minutes, and (4) Arena + Restraint: placed in one of the two chambers, allowed to explore for 5 minutes, and were then confined to a restraint tube for 25 minutes. Mice were then left undisturbed in their home cage for 50-60 minutes until perfusion.

### 2.4 c-Fos Immunohistochemistry

Mice were anesthetized with isoflurane and perfused transcardially with 1% PBS, followed by 4% paraformaldehyde. Brains were removed, post-fixed overnight at 4°C in 4% paraformaldehyde, and then transferred to 1% PBS. Brains were sectioned coronally (50 µm) using a vibratome (VT1000S, Leica Microsystems) and stored free-floating in 0.1M phosphate buffer at 4°C.

Immunohistochemistry was performed on every third slice throughout the rostral/caudal extent of the BLA, Hipp, and NAc. Free floating sections were incubated in blocking buffer (BB) (0.1 M phosphate buffer, 0.3% Triton X-100, and 0.5% normal goat serum) for 2 h at room temperature, followed by rabbit anti-c-Fos (abcam ab190289; 1:5,000 in BB) overnight at 4 °C. Sections were washed 3x with rinse buffer (0.1 M phosphate buffer, 0.3% Triton X-100) and incubated with goat anti-rabbit 488 (Alexa Fluor 488 (Invitrogen A11008; 1:500 in BB) for 2 hours at room temperature. After washing 3x with rinse buffer, sections were stored in 0.1 M phosphate buffer until mounting. Sections were mounted and coverslipped with Vectashield mounting media containing DAPI.

### 2.5 Imaging and Analysis

Images were captured using a Zeiss Axio Zoom V.16 microscope at 112x magnification. c-Fos+ cells were quantified from BLA, NAc, and dorsal and ventral hippocampal CA1 using ImageJ. Cell counts were normalized to the measured area of each region of interest, and these values were compared between the four groups described in 2.3. Some mice were eliminated from analysis due to inadequate sectioning or IHC processing.

### 2.6 Statistical analysis

All statistical analyses were performed using GraphPad Prism 10.3.0. Data were analyzed using a two-way ANOVA with Sidak’s multiple comparisons test. Data are presented as a box and whiskers plot. The line in the middle of the box represents the median. The box extends from the 25th to 75th percentiles. Whiskers represent minimum and maximum, and individual data points are shown. Schematics were produced in Biorender.

## 3 Results

### 3.1 Physical Restraint-Induced Conditioned Place Aversion

Mice were conditioned over several days in a two-chamber arena (Fig. 1a), and we compared the amount of time mice spent in the restraint-paired arena before and after conditioning. We found that mice spent significantly less time in the restraint-paired chamber after conditioning with no differences between males and females (Fig. 1b). This suggests that physical restraint effectively induces CPA similarly in both sexes.

### 3.2 c-Fos Expression Analysis Reveals Region-specific Activation Patterns

To determine which brain regions are engaged when an aversive stimulus like restraint is associated with contextual features of the environment, we examined c-Fos expression after mice were exposed to one of four conditions: undisturbed home cage housing, restraint stress in the home cage, exposure to a CPA chamber alone, and restraint stress in one of the chambers of the CPA arena (Fig. 2a). The latter two conditions mimic the CPA conditioning sessions described above. We quantified c-Fos+ cells in the BLA, NAc, and Hipp, three areas implicated in stress and learning (Fig. 2b).

**Figure 2.**
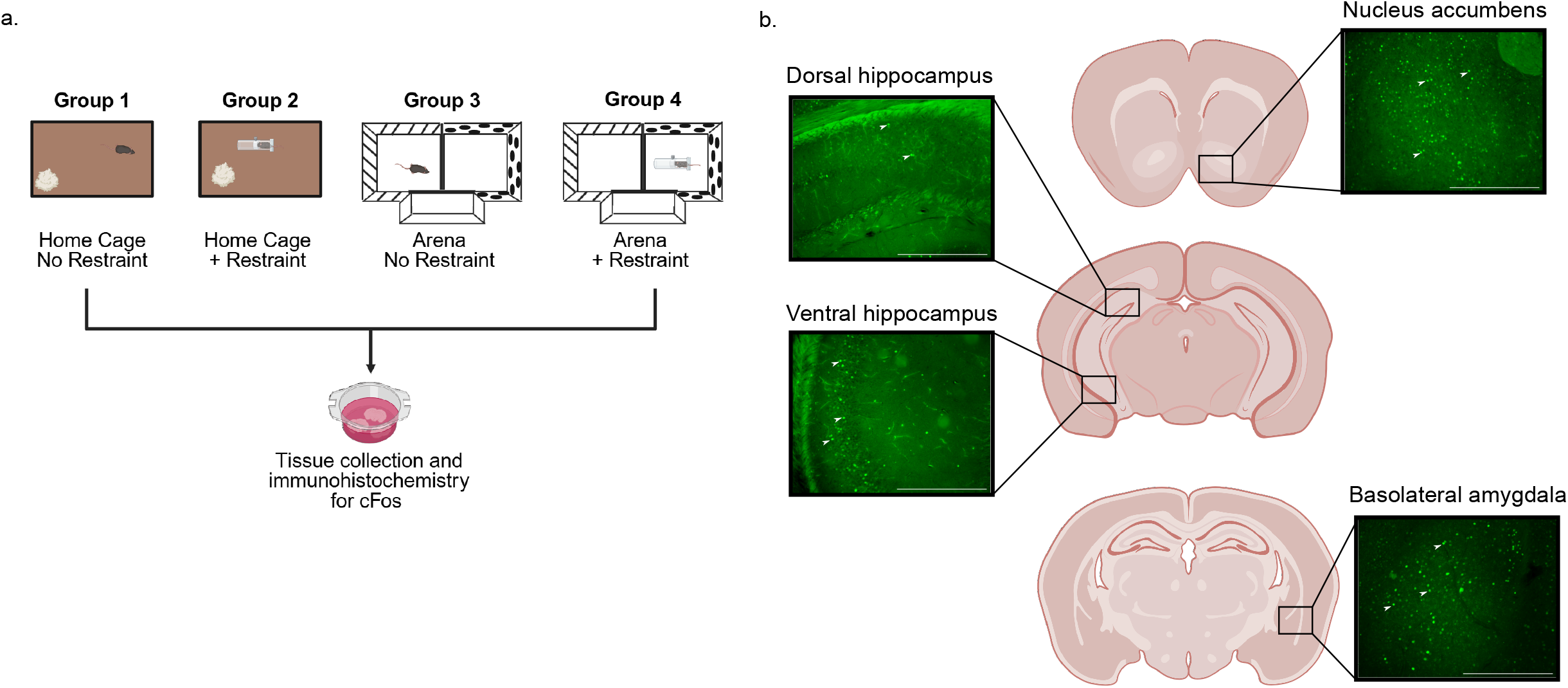
Determining areas of activation during conditioning using cFos expression. **a)** Experiment schematic. **b)** Representative images of the NAc, BLA, dHipp, and vHipp. Arrows indicate c-Fos+ cells. Scalebars represent 100µm.

In the NAc and BLA, we found that mice exposed to restraint in the CPA arena (restraint + arena group) showed a significant increase in c-Fos+ neurons (Fig. 3a-d). Neither restraint alone nor exposure to the arena alone produced significant activation in these regions, suggesting preferential activation in response to stress and environmental novelty. Hippocampal activation appeared to be subregion specific. While we found no significant differences in c-Fos expression in dorsal CA1 (Fig 3e-f), arena exposure led to a significant increase in c-Fos+ cells regardless of whether mice experienced restraint (Fig 3g-h). Together, these results suggest that the vHipp may be particularly sensitive to environmental novelty or the contextual cues in the chamber, whereas activation of the NAc and BLA activation involves both aversive and contextual stimuli.

**Figure 3.**
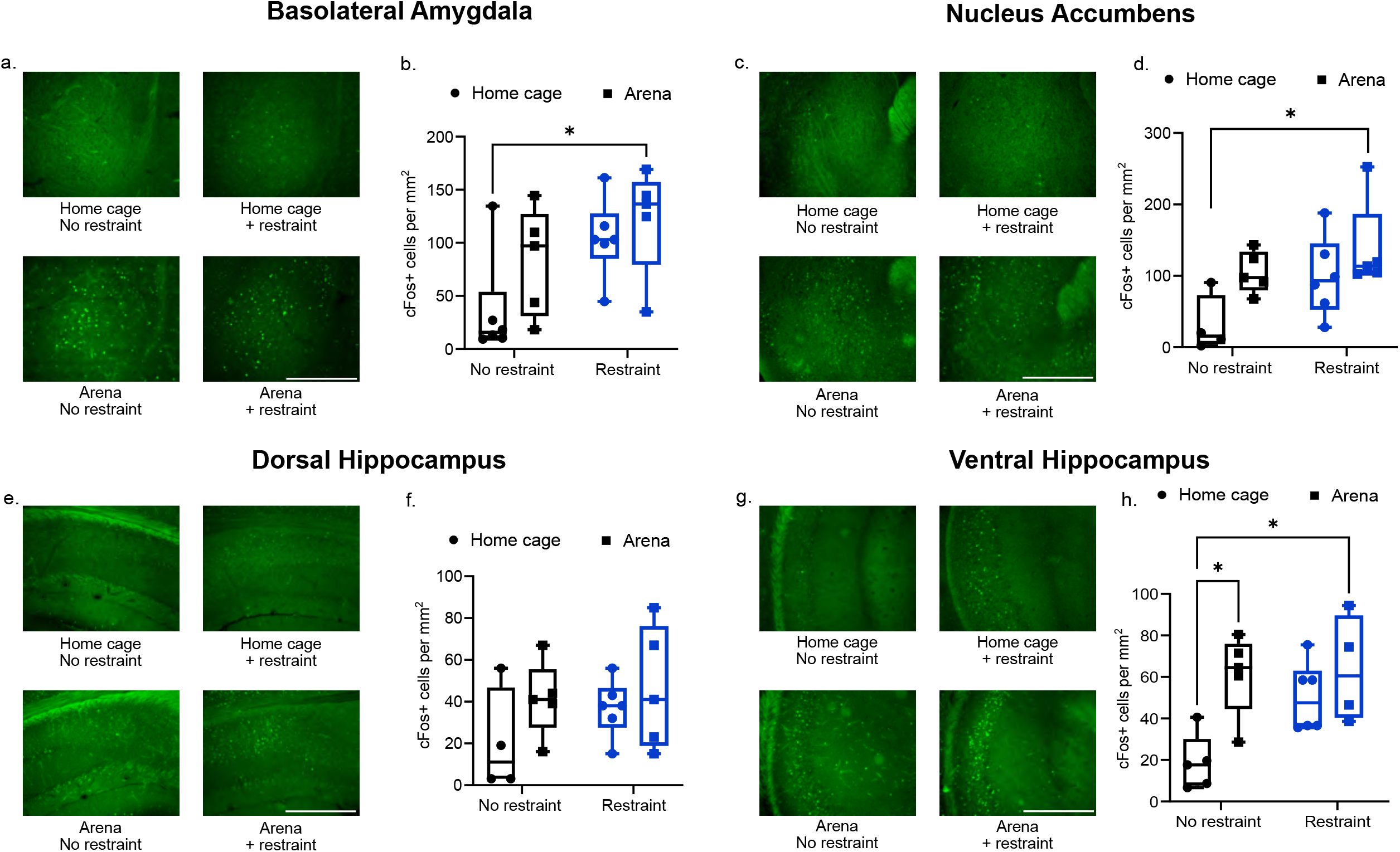
Distinct activation patterns in response to restraint stress and context. **a)** Representative images and **b)** quantification of c-Fos+ cells in BLA (n=6,5,6,6); Interaction: F_(1,18)_=0.5459, n^2^_p_ =0.0294; Main effect of restraint: F_(1,18)_=7.217, n^2^_p_ =0.286; Main effect of place: F_(1,18)_=2.576, n^2^_p_ =0.125; *p_(no restraint:home cage vs. restraint:arena)_=0.0420) **c)** Representative images and **d)** quantification of c-Fos+ cells from the NAc (n=4,5,6,5); Interaction: F_(1,16)_=0.5784, n^2^_p_ =0.0348; Main effect of restraint: F_(1,16)_=5.027, n^2^_p_ =0.239; Main effect of place: F_(1,16)_=6.254, n^2^_p_ =0.281; *p_(no restraint:home cage vs. restraint:arena)_=0.0334). **e)** Representative images and **f)** quantification of c-Fos+ cells from the dHipp (n=4,5,6,5); Interaction: F_(1,16)_=0.3695, n^2^_p_ =0.0226; Main effect of restraint: F_(1,16)_=1.201, n^2^_p_ =0.0699; Main effect of place: F_(1,16)_=2.383, n^2^_p_ =0.130). **g)** Representative images and **h)** quantification of c-Fos+ cells from the vHipp (n=5,5,6,4); Interaction: F_(1,16)_=2.968, n^2^_p_ =0.156; Main effect of restraint: F_(1,16)_ =4.021, n^2^_p_ =0.201; Main effect of place: F_(1,16)_ =10.84, n^2^_p_ =0.404; *p_(no restraint:home cage vs. no restraint:arena)_ =0.0148; p_(no restraint:home cage vs. restraint:arena)_ =0.0153). Scalebars represent 100µm.

## 4. Discussion

Stress-related psychiatric disorders, such as PTSD, are characterized by persistence and overgeneralization of learned associations between aversive events and the circumstances under which they occur [2]. Standard fear conditioning paradigms, invaluable for dissecting fear circuits, are limited in their ability to capture the active and persistent, context-driven avoidance that dominates presentation. Here, we use a restraint-induced CPA paradigm, to examine how stress-associated contexts drive avoidance behavior, providing a first step toward linking stress to the formation of persistent context-stress associations.

Our data demonstrate that repeated pairing of physical restraint with a contextually distinct chamber is sufficient to induce CPA, indicating that mice assign negative motivational value to restraint-paired context and subsequently avoid it when given a choice. Importantly, our paradigm isolates context-dependent avoidance from immediate escape or coping responses during the stress exposure itself, more closely analogous to the delayed, situational avoidance observed in response to adverse events. Restraint, a widely utilized stressor in rodents [9-12], also allows these findings to be integrated within the extensive literature on stress-induced neurobiological changes while incorporating an associative learning component.

Our c-Fos analysis revealed region-specific activation patterns. Exposure to the arena, with or without restraint, selectively increased c-Fos expression in the vHipp, in line with previously work demonstrating that this region is sensitive to environmental novelty and encodes emotionally-salient contextual representations [14-18]. In contrast, only the combination of restraint and contextual exposure produced robust c-Fos induction in the NAc and BLA, consistent with the well-described role for vHipp in conveying contextual and affective information to downstream regions (NAc and BLA) that integrate aversive motivational signals to guide approach-avoidance behavior [15,19-20]. This also has implications for determining the neuronal basis for overgeneralization and impaired extinction observed in PTSD and related disorders [2]. Overgeneralization of fear has been linked to disruptions in Hipp pattern separation and altered Hipp-amygdala connectivity [21-22]. Future studies employing circuit-specific manipulations and high-resolution approaches will be essential to test these hypotheses directly and determine how repeated stress reshapes activity, including distinguishing neuronal ensembles involved across learning phases (acquisition, consolidation, and retrieval).

Taken together, our work provides a tractable framework for probing how acute stress is transformed into context-bound aversive memories that drive avoidance. Placing restraint stress within the context of associative learning provides a foundation for understanding the neuronal mechanisms underlying the persistent, context-driven avoidance that is important for survival and altered in stress-related psychiatric disorders.

## Significance Statement

Our findings demonstrate restraint stress can induce persistent context-dependent avoidance and differential activation of stress-related circuits, establishing a novel model of adaptive avoidance.

## Acknowledgements

Microscopy was performed at the Keith Porter Imaging Facility in the College of Natural and Mathematical Sciences at UMBC. This work was supported by NSF IOS2402645, T32GM144876-02, the Meyerhoff Scholars program at UMBC, and start-up funds from UMBC. The authors report no conflicts of interest.

